# Development of a mass-spectrometry based method for the identification of the *in vivo* whole blood and plasma ADP-ribosylomes

**DOI:** 10.1101/2020.11.17.384719

**Authors:** Stephanie C. Lüthi, Anna Howald, Kathrin Nowak, Robert Graage, Giody Bartolomei, Christine Neupert, Xaver Sidler, Deena M. Leslie Pedrioli, Michael O. Hottiger

## Abstract

Blood and plasma proteins are heavily investigated as biomarkers for different diseases. However, the post-translational modification states of these proteins are rarely analyzed since blood contains many enzymes that rapidly remove these modification after sampling. In contrast to the well-described role of protein ADP-ribosylation in cells and organs, its role in blood remains mostly uncharacterized. Here, we discovered that plasma phosphodiesterases and/or ADP-ribosylhydrolases rapidly demodify in vitro ADP-ribosylated proteins. Thus, to identify the in vivo whole blood and plasma ADP-ribosylomes, we established a novel mass-spectrometry based workflow that was applied to blood samples collected from LPS-treated pigs (Sus scrofa), which serves as a model for human systemic inflammatory response syndrome. These analyses identified 60 ADP-ribosylated proteins, 17 of which were ADP-ribosylated plasma proteins. This new protocol provides an important step forward for the rapidly developing field of ADP-ribosylation and defines the blood and plasma ADP-ribosylomes under both healthy and disease conditions.

## INTRODUCTION

The clinical condition of systemic inflammatory response syndrome (SIRS) or sepsis is caused by pathogen-associated molecular pattern (PAMP) molecules, including lipopolysaccharide (LPS)^1^. These molecules interact with toll-like receptors (TLRs) on host immune cells, which triggers increased production of pro-inflammatory cytokines by immune cells circulating in the blood^2, 3^. Studies suggest that during sepsis, control of the primarily physiological host response to the infection is lost, which in turn causes a multitude of systemic dysfunctions that ultimately lead to multi-organ failure and the resulting high mortality rates^4^. Although sepsis has been extensively studied, the specific molecular mechanisms that drive multi-organ failure and eventual death are not fully understood^4, 5^. As sepsis is one of the leading causes of death in humans, developing a reproducible study system and accurate diagnostic tools that allow sepsis detection at early stages is critical to facilitate appropriate therapeutic interventions and developments^5^. Porcine pleuropneumonia, a highly infectious disease in pigs that is caused by *Actinobacillus pleuropneumoniae*, could serve as a model for human SIRS and sepsis^6^. LPS derived from *A. pleuropneumoniae* has been shown to induce inflammatory responses by triggering alveolar macrophages to overproduce proinflammatory cytokines like IL-1B, IL-6 and TNFa^7^. This proinflammatory cytokine influx, combined with the occurrence of typical clinical symptoms such as fever and elevated heart and respiratory rates match the defined criteria of human SIRS^3, 6^. Studies highlighting the similarity between the proteomes of domestic pigs (*Sus scrofa domesticus*) and humans further emphasized the potential of pigs as model systems for human patients, especially for infectious disease and inflammation research^8^.

Plasma is a readily accessible matrix for protein and biomarker research, and the plasma proteome has been extensively studied in humans, and various animals including pigs, under both healthy conditions and during the acute phase reaction; which is a complex early defense mechanism that the body mounts in response to a triggering event like infection or inflammation^9, 10^. Plasma proteins whose abundances increase in response to inflammation or tissue damage have been termed positive acute phase proteins (pAPPs). Modern medicine relies on the detection of these pAPPs to distinguish between healthy and diseased human patients on an individual basis, as well as large scale monitoring of herd health in farm animals^10^. However, the abundance of plasma proteins is not the only relevant feature; their modification states have also been previously studied as protein function can be regulated by post-translational modifications (PTMs)^11^. These studies have led to the discovery of glycosylated proteins as prognostic markers for certain cancer types and phosphorylated proteins as prognostic markers for various diseases^12, 13^. The role of other PTMs in plasma have not yet been examined.

ADP-ribosylation is a dynamic and reversible PTM that transfers one (MARylation) or multiple (PARylation) ADP-ribose moieties from NAD^+^ to an amino acid acceptor site of a target protein^14^. The possible acceptor amino acids include lysine, arginine, aspartic acid, glutamic acid, cysteine, and serine, as well as the recently discovered tyrosine and histidine^15–20^. Different ADP-ribosyltransferases (ARTs) have been described to MARylate or PARylate proteins. These enzymes are grouped into three subgroups based on their structural similarities to either diphtheria-toxin (ARTDs), cholera-toxin (ARTCs) or sirtuins^21^. Other enzymes are able to either hydrolyze the PAR chains (i.e. PARG), leaving a single ADP-ribose moiety bound to the protein, or erase the modification completely^22^. ADP-ribosylation plays a crucial role in many cellular processes including regulation of cellular inflammatory responses^14^. In contrast to the well-described role of ADP-ribosylation in such cellular mechanisms, its role in blood remains mostly uncharacterized. So far there has only been a single study reporting a correlation between non-enzymatic ADP-ribosylated serum proteins and cancer^23^, but studies have not yet linked blood and/or plasma protein ADP-ribosylation to inflammatory diseases such as sepsis.

Here, we observed that serum prepared via standard methods identified very few ADP-ribosylated proteins by LC-MS/MS. This led us to first establish various *ex vivo* biochemical assays to determine whether ADP-ribosyltransferases and -hydrolases were present and active in the serum/plasma. We found that ADP-ribosylation could be induced by adding NAD^+^ to heat-denatured plasma samples and that erasers were also active in the blood. Importantly, these assays demonstrated that proteins are rapidly demodified after blood sample collection prior to MS sample preparation. Thus, standard blood sample collection techniques needed to be adapted to preserve the modification state of proteins at the exact timepoint of blood sample collection. To this end, we collected blood samples from control and LPS-treated pigs, both before and after treatment, and immediately denatured half of the blood sample with guanidine hydrochloride (GndHCl) to avoid demodification. The other half of the blood was used to prepare plasma. We applied an optimized ADPr-peptide enrichment workflow, that we established here, to the denatured porcine blood samples to define the *in vivo* blood and plasma ADP-ribosylomes and determine how modification states differed between healthy and LPS-treated pigs. Interestingly, histidine was identified as one of the main ADPr-acceptor amino acids in blood and plasma samples, suggesting that the main writer of ADP-ribosylation in the blood has yet to be identified.

## MATERIALS AND METHODS

### LPS-induced SIRS study system

This study was conducted in accordance with the Swiss animal protection ordinance, under animal experimentation license number ZH046/17. 4-6-week-old pigs (n=10) were housed at the University of Zurich animal hospital and randomly allocated to either the control (n=5) or LPS group (n=5). The LPS trial lasted for a total of 10 days. From day 1 to 8 pigs were allowed to adapt to the new housing facilities and their clinical conditions assessed daily based on dyspnea, coughing, lung auscultation, nasal discharge and behavior parameters, which allowed us to define their health status and determine if individual animals should be removed from the trial. None of the pigs entered into the study showed signs of porcine pleuropneumonia, thus no animals were removed from the trial. On day 9 all pigs were again examined clinically, weighed, and a first blood sample collected. Pigs were then injected intramuscularly with 0.9% NaCl-solution (control group, 250 μl) or with LPS derived from *A. pleuropneumoniae* (treatment group, 250 μl 25% Montanide 25 containing 31.3 μg LPS). Following the injections, core body temperatures were assessed after 4, 6, 8, and 10 hrs and a second blood sample was collected from each pig at the 8 hrs timepoint.

### Blood sample collection and sample preparation

6 mL of venous blood were collected from each pig at both timepoints as previously described^24^. After collection, samples were immediately split into 3 fractions: 2 ml were transferred to heparin and EDTA tubes respectively (both BD, Franklin Lakes, New Jersey). Heparin samples were then centrifuged (1000x g, 10 min), the plasma separated from the cell pellet and stored at −80 °C for downstream plasma proteome analyses. The EDTA samples were kept at RT and immediately processed for hematological analyses. The remaining 2 ml were added to falcon tubes containing 4 ml of 6 M GndHCl (Sigma Aldrich, Saint Louis, Missouri) and mixed thoroughly to denature the samples. These whole blood lysates (WBL) were then heated to 95 °C for 10 min and stored at −80 °C until LC-MS/MS ADP-ribosylome analysis.

### Western blot analysis

For Western blot (WB) analyses, samples were prepared using standard SDS-PAGE and blotting techniques. Membranes were incubated with blocking solution (3% BSA in TBS-T (0.15 M NaCl, 10 mM Tris pH 7.5, 0.05% Tween-20)) for 1-2 hrs at room temperature (RT), and subsequently incubated overnight at 4 °C with the primary anti-ADPr-antibody (diluted 1:10’000 in TBS-T; Cell Signaling Technology, Leiden, Netherlands^25, 26^). Membranes were washed 3x for 5 min with TBS-T and incubated IRDye goat-anti-rabbit IgG secondary antibody (diluted 1:15’000 in TBS-T, LI-COR Biosciences, Bad Homburg, Germany) for 1 hr at RT and washed again as described before scanning (Odyssey Scanner, LI-COR Biosciences, Bad Homburg, Germany).

### Auto-modification of ARTD1

ARTD1 (formerly PARP1, Trevigen, Gaithersburg, Maryland) was auto-modified by incubating 10 U ARTD1 with or without 4.5 μg HPF1, 2 μg activated DNA and 5 μM NAD^+^ in 1x Trevigen PARP1 reaction buffer (Trevigen, Gaithersburg, Maryland) at 25 °C for 20 min^19^. The reaction was stopped by adding Laemmli buffer (final concentration 1x) and heating to 95 °C for 10 min. Alternatively, for the experiment characterizing eraser activity in plasma, ARTD1 (in-house) was auto-modified by incubating 0.875 μg ARTD1 with or without 100 μM NAD^+^, activated DNA (1 μg/reaction) in 1x RB (50 mM Tris-HCl pH 7.4, 4 mM MgCl2, 250 μM DTT) at 37 °C for 1 hr (total volume of 25 μl per reaction).

### Induction of ADP-ribosylation *ex vivo* via heat denaturing

25 μl of porcine plasma were heat denatured (95 °C, 10 min) in presence or without 100 μM NAD^+^ and subsequently incubated at 37 °C overnight to induce ADP-ribosylation. Samples were then diluted with 50 mM Tris pH 7.4 and 6x Laemmli buffer was added (final concentration 1x), sonicated and heated (95 °C, 10 min) before storing at −20 °C. GndHCl containing samples were subjected to a buffer exchange regime using Microcon-10 kDa centrifugal filter units (Merck, Darmstadt, Germany). Centrifugation (14’000x g, 20 min, RT) and washing 2 times with 100 μL 50 mM Tris pH 7.4 led to buffer exchange and SDS-PAGE compatibility. The resulting samples were resuspended in 25 μL Tris pH 7.4 and prepared as described above.

### Induction of ADP-ribosylation *ex vivo* via hARTC1

DC27.10 cells expressing human ARTC1 (hARTC1) on the cell membrane (kindly provided by F. Koch-Nolte) were grown in RPMI medium (1x) with GlutaMax supplement (Thermo Fisher Scientific Inc., Waltham, Massachusetts) and with 5% fetal calf serum at 37 °C. 2 x 10^6^ cells were collected via centrifugation (300x g, 5 min, 4 °C) and gently resuspended in 100 μl of plasma. Then, 100 μM NAD^+^ was added to the cell suspension and samples were incubated at 37 °C for 1 hr. The resulting plasma was then collected by centrifugation (300x g, 5 min, 4 °C) and analyzed via WB.

### Cell culture and ADPr-peptide enrichment

HeLa cells were cultured and treated with H_2_O_2_ as previously described and the lysates were used to test the new workflow and served as positive and negative controls for LC-MS/MS of the WBLs^16^. Wildtype and an evolved Af152 macrodomain, with 1000-fold enhanced binding affinity to ADP-ribose (*Nowak et al*., Nat.Com. manuscript accepted), were expressed and purified as previously described^27^. Binding and crosslinking of the GST-Af1521 fusion proteins to Glutathione Sepharose 4B beads (Sigma Aldrich, Saint Louis, Missouri), peptide enrichment (“standard workflow”) and pre-MS clean-up using stage tips were performed as previously described^27^”^29^. After elution, samples were resuspended to 12 μl in MS-buffer (3% ACN, 0.1% formic acid), vortexed briefly, sonicated (10 min, 100%, 25 °C), and centrifuged (16’000x g, 2 min, RT) before MS analysis.

For the new workflow established in this study, samples were diluted to 0.2 M GndHCl using 1x PARG buffer (50 mM Tris-HCl pH 8.0, 50 mM NaCl, 10 mM MgCl2, 0.25 mM DTT), and 4.2 μg PARG (in-house) per 10 mg protein and 10 μl of Benzonase (Merck Millipore, Darmstadt, Germany) per 30 ml were then added to each sample. The samples were incubated at 37 °C for 1 hr. Samples were then trypsin digested according to standard protocol and all subsequent steps performed as previously described. WBL samples collected during the LPS trial were prepared using the new workflow with additional high pH (HpH) and low pH (LpH) fractionation on MicroSpin C18 columns (Nest Group Inc., Southborough, Massachusetts) as described in^20^. Samples were eluted from the HpH column using three different percentages of ACN (7%, 15%, and 50% in 20 mM NH4OH) and from the LpH column using one condition containing 60% ACN / 0.1% TFA.

### Preparation of plasma input samples for MS analyses

Plasma input samples were prepared via filter aided sample preparation (FASP)^30^. Briefly, samples were denatured using GndHCl lysis buffer (6 M GndHCl, 5 mM TCEP, 10 mM CAA, 100 mM Tris-HCl pH 8.0, final concentration 4 M), heated (10 min at 99 °C) and sonicated. 100 μg of protein were transferred onto the Microcon-10 kDa centrifugal filter unit, centrifuged (14’000x g, 20 min, RT) and washed 3 times with 100 μl 0.5 M NaCl (centrifugation as described). 120 μl of 50 mM ammonium bicarbonate were added to the filter unit and centrifuged as described above. Trypsin (Promega, Fitchburg, Wisconsin) was added at a 1 μg trypsin : 25 μg protein ratio, the filter units sealed and incubated overnight (37 °C, 16-20 hrs, 300 rpm). The filter units were then centrifuged as described and the columns re-eluted using 80 μl 50 mM ammonium bicarbonate solution. The eluate was acidified with 10% TFA to a pH of 3-4 and the samples were then further prepared for MS analysis via Zip Tip clean-up according to the Millipore zip tip user guide (Merck Millipore, Darmstadt, Germany). Samples were resuspended in MS-buffer and processed as described for ADPr samples.

### Mass spectrometry and data analysis

LC-MS/MS analyses of ADP-ribosylated peptides were performed using the Product Preview method as described in Bilan *et al*. at the Functional Genomics Center Zurich using the Orbitrap Fusion Lumos mass spectrometer (Thermo Fisher Scientific Inc., Waltham, Massachusetts)^31^. Briefly, 2-5 μl of the prepared peptides solutions were eluted over 112 min at 300 nl/min, with ACN concentrations ranging from 5% to 95%. Initial high energy HCD scans were then applied to detect unique fragmentation ions of ADP-ribose, which in turn triggered product-dependent MS-events. Parallel high resolution HCD and EThcD scans were acquired to maximize peptide sequence coverage and PTM localization confidences^31^.

Downstream data analysis was performed as previously described ^31^. The HCD and EThcD deconvoluted separated .msg files were searched against the UniProt database for human and porcine proteins respectively using Mascot, and the following search parameters were applied: trypsin digests with up to 5 missed cleavages, carbamidomethyl as a fixed modification on cysteine (C), acetyl (protein N-term), oxidation (M), and ADPr_HCD_249_347_583 DEHKRSY (HCD scans, neutral losses defined in ^32^ and *Gehrig et al*., manuscript under revision) or ADP-Ribosyl EThcD (for EThcD scans) as variable modifications, peptide tolerance =10 ppm, number of ^13^C = 1, peptide charge = 2^+^/3^+^/4^+^, MS/MS tolerance = 0.05 Da^33^. Briefly, the resulting files were filtered to include only peptides with one or more ADP-ribose modifications, peptide scores > 15 (WBL samples) or > 20 (HeLa cell lysates) and peptide expect values < 0.05. For ADP-ribose acceptor site localizations, the list of modified peptides was further filtered based on peptide variable modification confidence values > 95%. MS analyses of input proteome samples were performed using standard data dependent acquisition (DDA) methods. The following search parameters were used to search input proteome deconvoluted data: trypsin digests with up to 5 missed cleavages, carbamidomethyl as a fixed modification on cysteine (C), acetyl (protein N-term), oxidation (M), peptide tolerance =10 ppm, number of ^13^C = 1, peptide charge = 2^+^/3^+^/4^+^, MS/MS tolerance = 0.5 Da^33^. Plasma proteome search results were filtered as described above for peptide score and peptides expect values, without regard to ADP-ribosylation. The mass spectrometry proteomics data have been deposited to the ProteomeXchange Consortium via the PRIDE partner repository with the dataset identifier PXD022156.

### Statistical analysis

Statistical analyses were performed using R version 3.5.2^34^. Differing core body temperatures and total leukocyte amounts of control/LPS-treated pigs were tested for their statistical significance at each time point (0 hr, 4 hrs, 6 hrs, 8 hrs and 10 hrs post injection or before and after injection, respectively). Normal distribution was first tested using a ShapiroWilk normality test and subsequently a two-tailed, unpaired t-test (normal distribution) or a MannWhitney U test (non-normal distribution) was applied. Differences in peptide presence were analyzed by creating 2 x 2 contingency tables for each ADP-ribosylated peptide found in plasma samples and then categorizing the data as either (1) control conditions or LPS treatment and (2) number of pigs where the peptide was detected or not detected. Due to a non-normal distribution as well as unequal and small sample sizes a Fisher’s exact test was used for further analysis. All results were considered significant for p-values < 0.05.

## RESULTS

### LPS treatment significantly increased core body temperature and total leukocyte numbers in pigs

The LPS-induced SIRS study system consisted of an LPS trial during which 10 pigs were injected with either NaCl (n=5) or LPS (n=5) solutions (Suppl. Table 1). Blood samples were collected at different timepoints. To determine how the animals reacted to a single LPS injection (31.3 μg), the core body temperature and total number of leukocytes were analyzed before and after injection, as these are two key symptoms of SIRS. The core body temperatures of the LPS-treated pigs increased significantly 4, 6, 8, and 10 hrs after injection compared to the control group (Figure 1A). The total amount of white blood cells also increased significantly in the LPS treatment group after injection, while significant changes were not observed before or after NaCl injection in the control group (Figure 1B). Finally, protein concentrations were defined for each plasma sample taken before and after the LPS or NaCl injections. Significant differences were not observed between or within the two groups at either timepoint (Figure 1C). Together, these data indicate that the established porcine LPS-induced SIRS model elicits clinical and hematological reactions similar to human SIRS.

**Figure 1:**
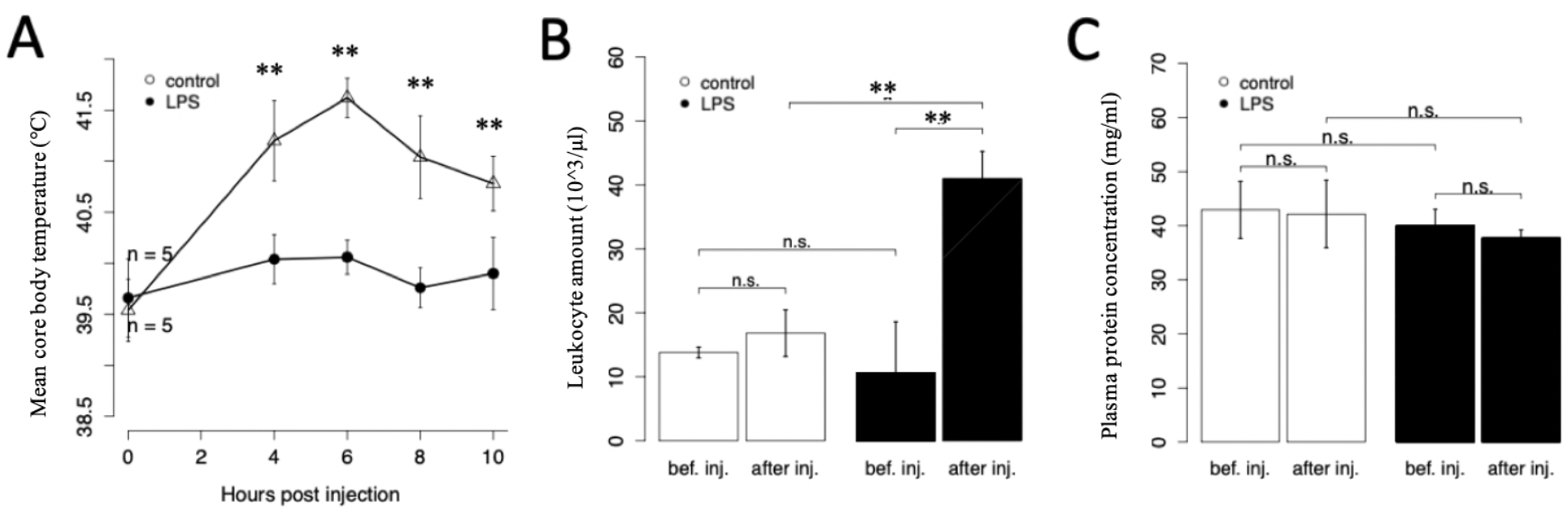
LPS injections triggered significant increases in core body temperature and blood leukocyte concentrations. A: Mean core body temperature (°C) of pigs ± standard deviation after NaCl (control group) or LPS (treatment group) injections at indicated timepoints during the LPS trial. B: Mean leukocyte amounts (x 10^3^/μl) ± standard deviation before and after NaCl (control group) or LPS (treatment group) injections. C: Mean plasma protein concentration ± standard deviation before (bef. inj.) and after (after inj.) NaCl (control group) or LPS (treatment group) injections. Significant differences are labeled with an asterisk (* for p<0.05 – p=0.01, ** for p<0.01 - p=0.001, *** for p<0.001), while non-significant differences are labeled “n.s.”.

### Optimization of the MS-based workflow to allow detection of low abundant ADP-ribosylated proteins

Preliminary Western Blot (WB) and MS-based experiments with serum samples from NaCl- and LPS-treated pigs demonstrated that the levels of ADP-ribosylation in these samples were very low, even below the level of untreated HeLa cell culture samples (data not shown). We, therefore, first optimized the standard ADP-ribosylome workflow using H_2_O_2_-treated HeLa cells to increase enrichment efficiency prior to MS-based ADP-ribosylome analyses ^16, 32^. Here, we included a major change to the established standard workflow; PARG/Benzonase lysate treatments were moved to the first step of the sample preparation protocol in the hope that this would improve trypsin digestion efficiency and modified peptide enrichment (Suppl. Figure 1). This was rationalized based on recent studies demonstrating that S-ADPr modifications are typically found within KS and RS motifs^31, 35^, thus reducing PAR chains to MAR modifications prior to trypsin digestion could improve protein-to-peptide digestion. In addition, we also reasoned that C18 column clean-up after PARG digested samples would eliminate most of the free ADP-ribose prior to ADPr-peptide enrichment. This could provide an additional advantage, as competitive binding between the free ADP-ribose and ADPr-modified peptides to the Af1521 macrodomains would be reduced and, thus, MARylated peptide enrichment efficiencies should increase.

To determine if this protocol modification improved ADP-ribosylome coverage, we compared the unique peptides, unique modification sites (> 95% localization confidence) and unique modified proteins identified for the standard workflow and new workflow (Figure 2A). The new workflow outperformed the standard workflow considerably by identifying ~3x more modified peptides, ~2x more unique modification sites and ~2.5x more proteins, suggesting that this modification provided an experimental advantage (Figure 2A, Suppl. Table 2). Comparison of the standard workflow and new workflow coverages revealed that only 5 proteins (5.4%) were uniquely identified in samples using the standard workflow, indicating that the sample preparation changes made did not bias peptide enrichment or modified protein identifications (Figure 2B). Importantly, the new workflow identified 59 proteins that were not detected with the standard workflow and greatly reduced the number of mis-cleaved peptides (Figure 2C). In an attempt to further increase our ADP-ribosylome coverage, we applied additional sample preparation improvements that were recently established^20^. Here, lower acetonitrile concentrations were used when peptides were eluted from the SepPak columns during the first clean-up step and the high pH and low pH Stage-tip fractionation protocol was applied during the final clean-up to reduce sample complexity and improve peptide sequencing during MS analysis. Applying these modifications, in conjunction with the workflow modifications described above, further increased ADP-ribosylated peptide, modification site and protein detection rates by 68.7%, 52.9%, and 69%, respectively (Suppl. Figure 2A & B, Suppl. Table 3). Taken together, these data demonstrate that these simple sample preparation modifications increased ADP-ribosylome coverage considerably and were used for MS sample preparation of the LPS trial derived blood samples.

**Figure 2:**
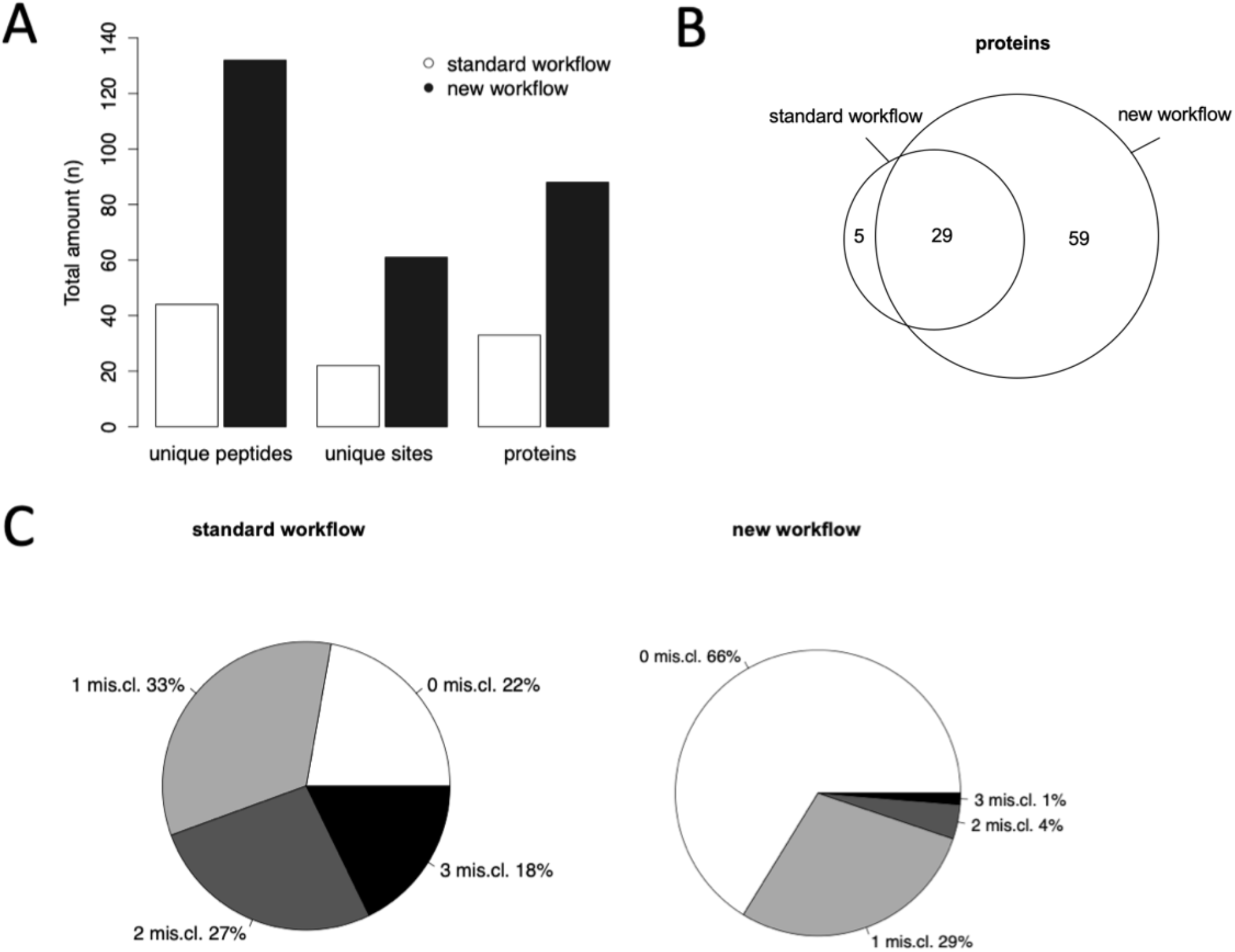
Comparison of the standard and new ADP-ribosylome MS sample preparation workflows using HeLa H_2_O_2_-treated cell lysates. A: Comparison of the number of unique peptides, unique ADPr-modification sites (>95% localization confidence) and unique modified proteins identified in HeLa H_2_O_2_-treated cell lysates by LC-MS/MS analyses when samples were prepared using either the standard or new workflow. Combined HCD and EThcD data are shown. B: Venn diagrams showing the overlap of ADP-ribosylated proteins that were identified using either the standard or new workflow. C: Comparison of the amounts of mis-cleaved (mis.cl.) peptides from HeLa H_2_O_2_-treated cell lysates prepared for MS analysis using the standard workflow (left panel) or new workflow (right panel).

### Endogenous erasers of ADP-ribosylation are active in blood samples and rapidly reduce ADP-ribosylation after sample collection

Despite the sample preparation improvements described above, very few ADP-ribosylated proteins were identified in serum samples collected from a preliminary LPS trial using this optimized ADP-ribosylome MS-workflow (data not shown). In an attempt to determine why this might be, we established various biochemical assays to establish if the ADP-ribosylation states of plasma/serum proteins could be altered during sample collection and preparation. Using a commercially available anti-pan-ADPr antibody^25, 26^, we wanted to determine whether plasma protein ADP-ribosylation could be rapidly erased after sample collection and/or during plasma preparation. To this end, ADP-ribosylation was induced in healthy pig plasma samples *ex vivo* via incubation with NAD^+^ and DC27.10 cells that ectopically express the R-ADPr-specific ADP-ribosyltransferase hARTC1^36^. After modification, the cells were removed, and the plasma incubated at 37 °C for 15 min, 1 hr, 4 hrs and overnight to determine if the ADP-ribosylation induced in the plasma could subsequently be erased. Indeed, a strong decrease of plasma protein ADP-ribosylation was observed already after 1 hr and 4 hrs (Figure 3A). Furthermore, ADP-ribosylation levels were reduced to those of untreated samples when the modified plasma samples were incubated overnight at 37 °C (Figure 3A), suggesting that R-ADPr modifications can be rapidly erased in plasma and blood samples.

**Figure 3:**
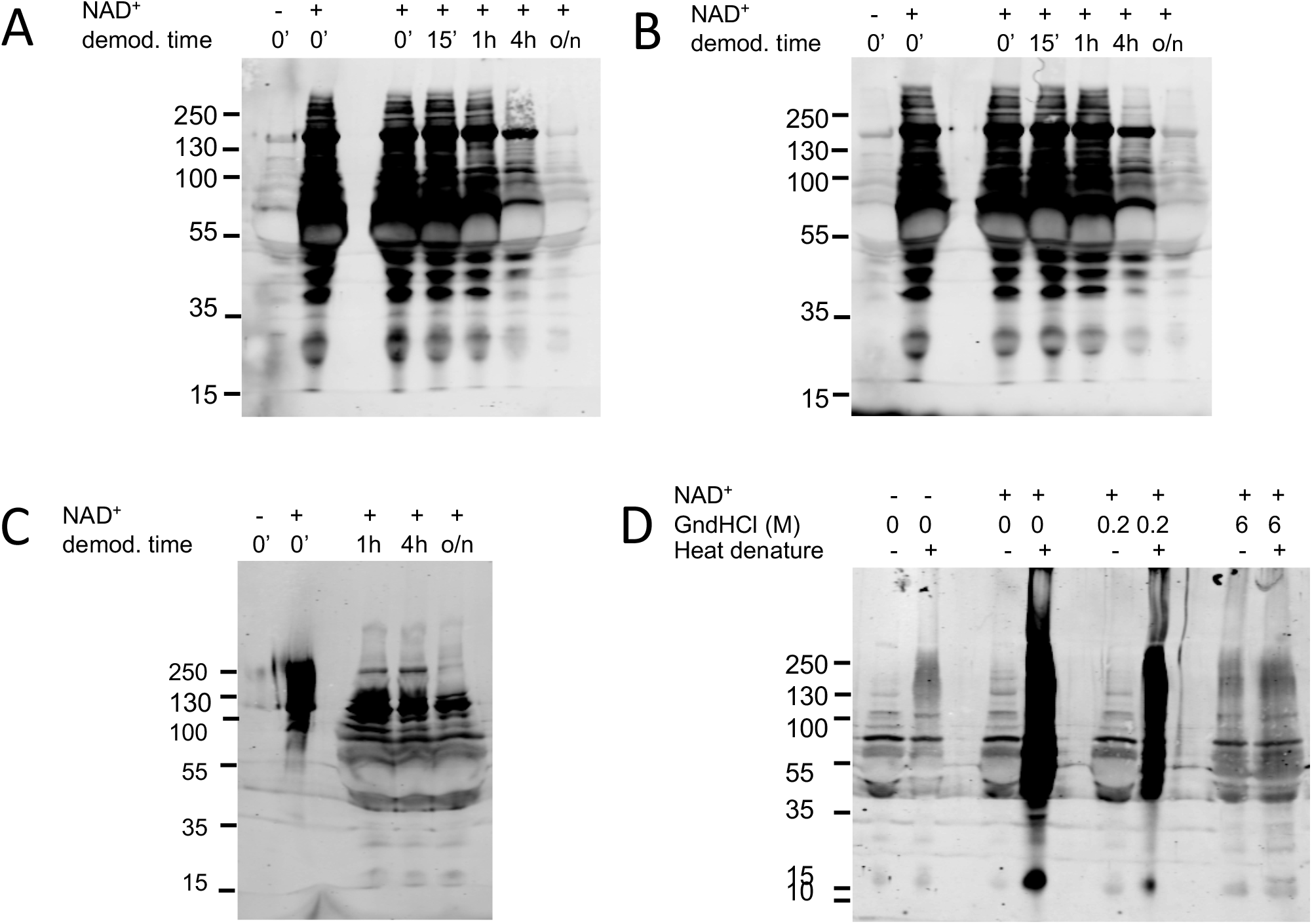
Writers and erasers of ADP-ribosylation are present and active in plasma. A: To determine how stable ADPr-modifications are in plasma samples, porcine plasma samples were ADP-ribosylated *ex vivo* via incubation with ARTC1 expressing DC27.10 cells and NAD^+^. The cells were removed and the plasma incubated for 0 min, 15 min, 1 hr, 4 hrs and overnight (o/n) at 37 °C. Western blot analysis of the plasma using the Cell Signaling anti-ADPr-antibody were then used to determine if ADP-ribosylation levels decreased during postmodification incubation (demod. time). B: To determine if the observed changes were enzymatically driven, 100 μM cAMP was added to the samples after removing the cells and the samples incubated as described above. C: ARTD1 was auto-modified and then incubated with porcine plasma from healthy pigs for 1 hr, 4 hrs, and overnight at 37 °C. Western blot analysis with the anti-ADPr-antibody was used to assess whether auto-modified ARTD1 could also be demodified by plasma. D: Western blot analysis of heat-denatured (95 °C, 10 min), GndHCl-denatured, and native porcine plasma using the Cell Signaling anti-ADPr-antibody.

Nevertheless, it remained unclear if the demodifications observed here were mediated by classic ADP-ribosylhydrolases that could be present in the blood or other enzymes, like phosphodiesterases (PDEs), that have been shown to be abundant in the blood and able to reduce ADP-ribose to phosphoribose^37, 38^. To investigate this, we repeated the experiment described above in the presence of cAMP, a metabolite that can compete with ADP-ribose as a PDE substrate. Interestingly, addition of 100 μM cAMP partially blocked the observed demodification for the first hour of incubation (Figure 3B). Together, these findings indicate that PDEs present in the plasma actively and rapidly erase mono-ADPr plasma protein modifications. These finding are critical, as blood collection and plasma preparation times fall within this time frame; thus, endogenous ADP-ribosylation modifications could be substantially reduced, or entirely lost, using standard sample preparation methodologies.

Having established that R-ADPr modifications could be erased in the plasma, we determined if this also holds true for other ADPr-amino acid acceptor modifications. To address this, we used ARTD1, which is mainly modified on serine (S) residues ^19, 39^. ARTD1 was automodified in the presence of NAD^+^, activated DNA and HPF1. Following auto-modification, ARTD1 was incubated with PJ34 and plasma 1 hr, 4 hrs, and overnight at 37 °C. WB analyses of these samples revealed that ARTD1 ADP-ribosylation states gradually decreased when plasma was present (Figure 3C). This was most evident following overnight incubation at 37 °C. Together, these findings suggest that ADP-ribosylhydrolases and PDEs are present in plasma samples and that *in vivo* ADP-ribosylated plasma proteins (regardless of ADPr-acceptor amino acid) can be demodified shortly after blood sampling prior to MS sample preparation.

### ADP-ribosyltransferases are present and active in porcine plasma samples

The presence and activity of ADP-ribosylation erasers in the blood prompted us to question whether ADP-ribosyltransferases are also present and active under ADP-ribosylome MS sample preparation conditions. To investigate this, plasma samples were heat-denatured or denatured with 6M or 0.2M guanidinium hydorchloride (GndHCl), and then incubated overnight at 37 °C in the presence or absence of NAD^+^. WB analyses revealed that extensive ADP-ribosylation was exclusively induced in plasma samples that were heat-denatured at 95 °C for 10 min and then incubated with exogenous NAD^+^ overnight at 37 °C (Figure 3D). Importantly, addition of GndHCl to the sample at a final concentration of 0.2M reduced the observed increase in ADP-ribosylation by about half and full denaturing conditions (6M GndHCl) inhibited ADP-ribosylation completely. Taken together, these studies demonstrate that both ADP-ribosyltransferases and ADP-ribosylation erasers (ADP-ribosylhydrolases and PDEs) are present and active in plasma samples. Thus, further optimization of the sample collection methods remains crucial to ensure that *in vivo* ADP-ribosylation states of these samples at the point of sample collection is preserved.

### ADP-ribosylome workflow sample preparation modifications allowed identification of the *in vivo* whole blood ADP-ribosylome

The findings presented above indicate that denaturing the samples completely with 6 M GndHCl immediately after blood collection could preserve the ADP-ribosylation state of the samples. To investigate this and identify the whole blood ADP-ribosylome of healthy and LPS-treated pigs via LC-MS/MS, we prepared whole blood lysates and matched plasma samples from the animals. The whole blood lysates (WBLs) were generated by denaturing the samples with 6 M GndHCl immediately after blood collection. At the same time, part of each blood sample was anti-coagulated and processed into plasma. The resulting WBLs were processed for LC-MS/MS analysis using the new ADP-ribosylome enrichment workflow with fractionation presented above and plasma samples were analyzed using standard shot gun LC-MS/MS methods to define the plasma proteomes of each of these samples (Suppl. Figure 3).

Following LC-MS/MS ADP-ribosylome analysis, we compared the number of unique peptides, unique ADPr-modification sites (> 95% localization confidence) and unique modified proteins identified in the WBLs of control and LPS-treated pigs. We found that similar numbers of ADP-ribosylated peptides and modified proteins were identified in each group, but ~1.5 x more unique modification sites (> 95% localization confidence) were identified in LPS-treated pigs compared to control pigs (Figure 4A). While a total of 60 ADP-ribosylated proteins were identified in the WBLs, we found that over two thirds of the proteins identified were unique to one treatment group or the other (Figure 4B and Suppl. Tables 4 and 5). Indeed, the control group contained 22 unique ADP-ribosylated proteins, while 21 modified proteins were specific to the LPS-treated group (Figure 4B). Given that most of the proteins were only identified in 1 of the 5 animals per treatment group (Suppl. Figures 4A and B), it is likely that these modifications are extremely low abundant. Interestingly, 17 ADP-ribosylated proteins were identified in both control and LPS-treated pigs, including the acute phase protein AHSG and hemoglobin subunits alpha and beta (Suppl. Figure 4C, Suppl. Tables 4 and 5). Statistically significant differences in the blood ADP-ribosylomes of control and LPS-treated groups were not observed on the protein level (likely due to small treatment group sizes (n=5)). Nevertheless, we did note that the identification rate of one of the HBA ADP-ribosylated peptides increased ~15% in the LPS-treatment group and that the identification rates of two HBB ADP-ribosylated peptides decreased >25%, while two other HBB ADPr-peptides recorded 75% and 35% increases in the identification rates in the LPS-treatment group (Suppl. Tables 4 and 5). Furthermore, the identification rates of AHSG, GAPDH and MDH1 decreased 40%, 75% and 40%, respectively, in the LPS-treated group.

**Figure 4:**
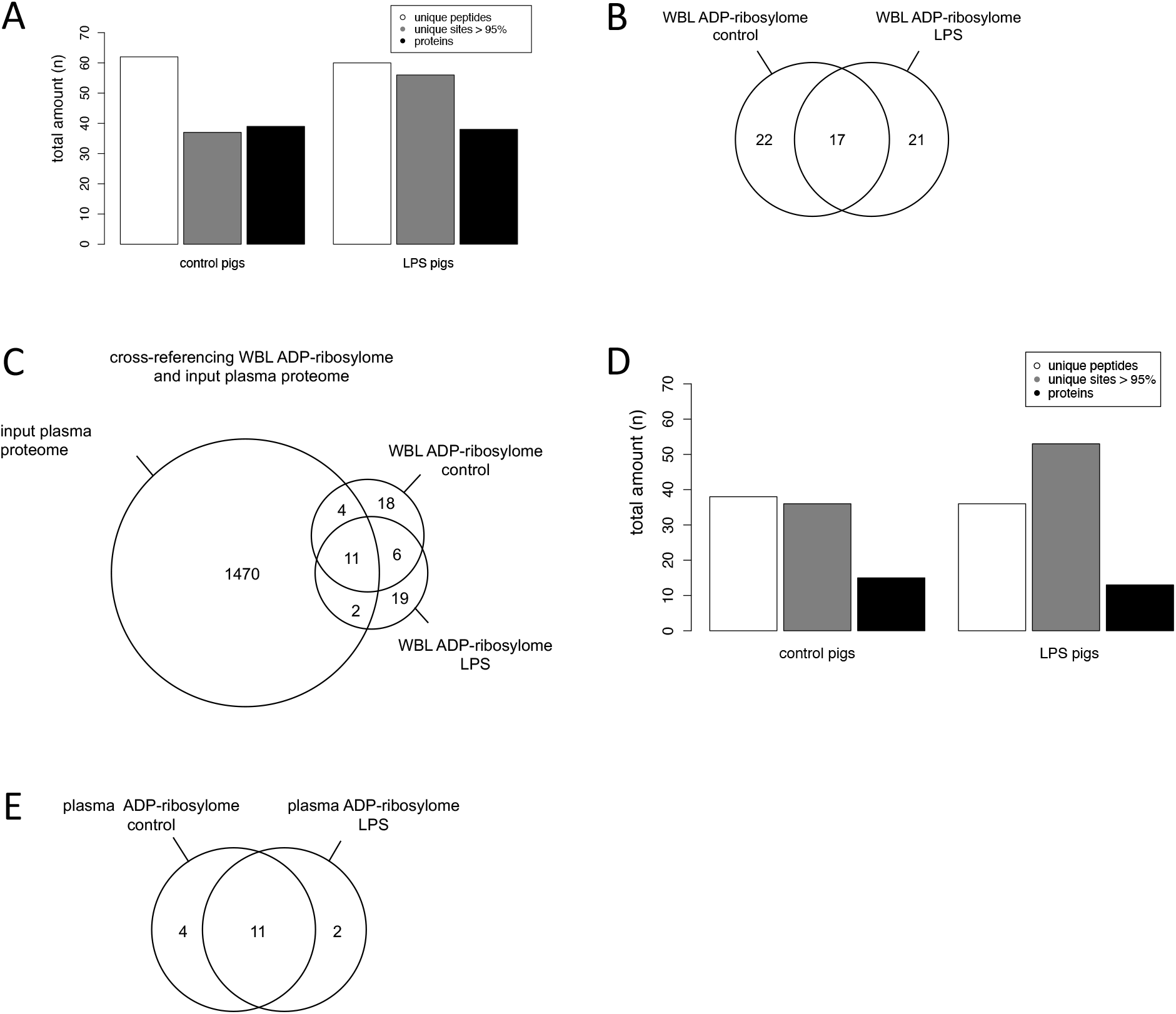
The *in vivo* WBL ADP-ribosylome identified in healthy and LPS-treated pigs. A: Overview of the unique ADP-ribosylated peptides, unique modification sites (>95% localization confidence) and proteins found in the WBL ADP-ribosylome of control and LPS-treated pigs. B: Venn diagram showing the overlap of WBL ADP-ribosylomes identified in healthy and LPS-treated pigs. C: The plasma ADP-ribosylome of control and LPS-treated pigs was determined by cross-referencing the WBL ADP-ribosylome of both groups with the reference plasma proteome (generated by combining the plasma proteomes of both groups). D: Overview of unique ADP-ribosylated peptides, unique modification sites (>95% localization confidence) and proteins found in the plasma ADP-ribosylome of control and LPS-treated pigs. E: Venn diagram showing the overlap of ADP-ribosylated plasma proteins identified in control and LPS-treated groups.

### Comparison of the WBL ADP-ribosylome with the corresponding plasma proteome defined the *in vivo* plasma ADP-ribosylome

We then set out to take this analysis further and separate the plasma ADP-ribosylome from the WBL ADP-ribosylome that is derived from cellular fraction of the blood. We rationalized that correlating the WBL ADP-ribosylome with the corresponding plasma proteome would allow us to define which of the WBL ADP-ribosylated proteins are plasma proteins and, thus, represent the plasma ADP-ribosylome (Suppl. Figure 3). To this extent, we defined the corresponding plasma proteomes of each WBL sample, which led to the identification of 1221 plasma proteins in healthy and 958 in LPS-treated pigs, respectively (Suppl. Table 6). To maximize plasma proteome coverage, these two plasma proteomes were combined to generate a pig plasma reference proteome, containing 1487 unique proteins, which were then compared to the WBL ADP-ribosylomes. This comparison identified the 17 ADP-ribosylated plasma proteins (Figure 4C). As observed with the WBL ADP-ribosylome, the control and LPS-treated pig plasma ADP-ribosylomes were very similar with respect to the number of unique peptides and ADP-ribosylated proteins identified (Figure 4D). Nevertheless, a handful of proteins were only identified in the control or LPS treatment groups (Figure 4E).

When analyzing protein ADP-ribosylation, the acceptor amino acids are of great interest since their distributions can help define the ADP-ribosyltransferases that could write the observed modifications. Thus, we determined the number of unique modification sites with > 95% PTM localization confidence in the WBL ADP-ribosylome, as well as the plasma ADP-ribosylome. Interestingly, H- and S-ADPr modifications were the most abundant ADPr-acceptor amino acids identified in both the WBL and plasma ADP-ribosylomes (Suppl. Tables 4 and 5). While the search algorithm also identified Y-ADPr, R-ADPr, D-ADPr and E-ADPr, H-ADPr and S-ADPr modification sites were ~2-3-times more abundant in both control and LPS-treated samples. Very similar distributions and amounts of unique acceptor sites with high PTM localization confidences were identified when analyzing the ADPr-acceptor sites of the plasma protein ADP-ribosylome. Together, these data demonstrate that the most prominent unique ADPr-acceptor sites with highest PTM localization confidence are indeed found within the plasma ADP-ribosylome and not the cellular fraction of the whole blood ADP-ribosylome. This could be due to the fact that plasma proteins, like HBB and AHSG are highly abundant and modified at multiple sites. Most interestingly, given that these results indicate that H-ADPr modifications are abundant here, it is likely that the most active ADP-ribosyltransferase in the plasma has not yet been characterized. In conclusion, applying the new workflow with fractionation to the WBL samples helped us identify the first blood ADP-ribosylome and the results indicate that there are most likely biologically relevant differences in the WBL and plasma ADP-ribosylomes of mock treated and LPS-treated pigs.

## DISCUSSION

In this study, we aimed to identify the whole blood and plasma protein ADP-ribosylomes in a porcine model for human SIRS and sepsis using our well-established MS-based ADP-ribosylome workflow^20, 31^. Unfortunately, preliminary experiments identified very few ADP-ribosylated proteins in serum samples from LPS-treated pigs. We, therefore, optimized ADP-ribosylated peptide enrichment and clean-up workflows that were used prior to LC-MS/MS analysis. While application of the sample preparation modifications to HeLa H_2_O_2_ lysates significantly increased ADP-ribosylated peptide and protein identifications compared to our current state-of-the-art ADP-ribosylome MS-workflow, it did not improve ADP-ribosylated protein identifications within the serum samples we collected from LPS-treated and healthy pigs. Surprisingly, using a number of *ex vivo* biochemical assays, we determined that ADP-ribosylhydrolases and/or PDEs and ADP-ribosyltransferases are present and active in plasma/serum. These finding indicated that proteins are rapidly demodified after blood sample collection prior to MS sample preparation, which prompted us to optimize the sample collection protocol to ensure that the modification states of the samples at that time would be maintained. To this end, we denatured blood samples from control and LPS-treated pigs immediately following collection and then defined the ADP-ribosylome of these whole blood lysates. Application of these protocol modifications allowed us to successfully define the first blood ADP-riboylomes, which contain 60 ADP-ribosylated blood proteins. To expand these analyses and define the corresponding plasma ADP-ribosylome, we cross-referenced the WBL ADP-ribosylomes with the total plasma proteome and identified 17 ADP-ribosylated plasma proteins. While differences between the control and SIRS/sepsis ADP-ribosylome samples were not significant in this study, the protocol optimizations presented here signify a significant advancement for the field of ADP-ribosylation. Indeed, the optimizations developed allowed us, for the first time, to define both the whole blood and plasma ADP-ribosylomes and, most importantly, can be applied to blood samples drawn from any species. Thus, this novel method is also applicable to human clinical samples and, as such, can support the identification of ADP-ribosylome specific biomarkers within a clinical setting.

The protocol optimizations that we present focused on improving ADPr-peptide enrichment efficiency and ADP-PSM identifications rate and quality. We demonstrated that i) reducing the long and/or branched PAR chains to MAR modifications before trypsin digestion improved the digestion site accessibility that significantly reduced protein miss-cleavage rates and ii) eliminating the free ADP-ribose produced following PARG PAR-to-MAR modification reduction significantly improved ADPr-peptide enrichments by reducing free ADP-ribose and the modified peptide Af1521 competitive binding dynamics. These conclusions are supported by recent studies demonstrating that oxidative stress induced S-ADPr modifications are typically found within KS and RS motifs^31, 35^. Thus, reducing the long S-ADPr PAR chains would allow more complete tryptic digestion, since trypsin is known to cleave peptide bonds after K and R^18, 40^. Finally, introducing the high pH/low pH fractionation steps at the end of the enrichment protocol further improved ADP-ribosylome coverage by reducing sample complexity, which is known to improve PSM identification rates and qualities^41^.

Subsequent definition of the plasma proteome led to the identification of ARTC3, a member of the ARTC ADP-ribosyltransferase protein family that is thought to be inactive due to alterations in the catalytic amino acid triad, in all control and LPS pig plasma samples. This ARTC has previously been identified in porcine muscle samples as well as human plasma samples ^42, 43^. Interestingly, one of the most abundantly modified amino acid identified here was histidine; one of the most recently identified ADPr-acceptor amino acid ^20^. This novel finding, together with the fact that ARTC3 is believed to be inactive, suggests that a low abundant and/or uncharacterized ADP-ribosyltransferase could be present in the blood and responsible for the modification of histidine residues on these target proteins. Previous studies identified sirtuins (SIRT1-5) and phosphodiesterases (PDE 5A & 12) in human plasma samples^42, 44^ and, indeed, we also identified PDE5A in porcine plasma. Since many known writers and erasers of ADP-ribosylation are localized within cells, they could be released into the blood after cell necrosis/lysis, and the resulting concentrations of writers and erasers in the plasma would most likely be very low, emphasizing the need for a highly sensitive method to detect these enzymes. Therefore, in depth analysis either via depletion and fractionation of the plasma proteome or parallel reaction monitoring (PRM)-MS could be applied to identify other writers or erasers that are present in plasma samples as previously described^44, 45^. Importantly, the presence of active ADP-ribosylation erasers in the blood/plasma suggests that other PTMs could also be erased during sample collection and plasma preparation. Thus, the blood/plasma sample preparation modifications that we developed here could prove beneficial when defining other PTMs, like phosphorylation or glycosylation, in plasma or serum.

Importantly, the novel method described here to define the blood and plasma ADP-ribosylomes provides an important step towards analyzing *in vivo* ADP-ribosylated blood and plasma proteins under clinical settings. The modified proteins that we have identified provide a first description of the *in vivo* whole blood and plasma ADP-ribosylomes. Interestingly, two of the most heavily modified proteins identified here in the WBL ADP-ribosylome were the hemoglobin alpha and beta subunits, which play a critical role in oxygen transport and are known to interact with various drugs^46^. In addition, we noted that the identification rates of some of the HBA, HBB, GAPDH and MDH1 ADP-ribosylated peptides varied between the control and LPS-treatment groups. We found that the number of HBA TYFPHFNLSHGSDQVK peptide PSMs increased by 15% in LPS-treated pigs. Interestingly, the ADPr-modification site for this peptide was localized to Y2, which is right next to the phenylalanine (F43) that interacts with heme group^47–49^; thus, it is possible that ADP-ribosylation of HBA at this position could interfere with hemoglobin function. Furthermore, we found that the corresponding peptide in HBB (FFESFGDLSNADAVMGNPK) was also ADP-ribosylated immediately next to F43 at E44 and that PSM identification rates for this peptide increased by ~40% in LPS-treated pigs. Lastly, we identified another HBB ADP-ribosylated peptide (VLQSFSDGLKHLDNLK) whose identification rate also increased by 40% and is fairly close to H63 - a residue that is conserved across most species and is directly involved in the oxygen binding site^47, 49^. It is intriguing to speculate how hemoglobin subunit ADP-ribosylation could alter oxygen, carbon dioxide, carbon monoxide and/or drug binding affinities; could the modification alter binding in a manner similar to what has been described for different hemoglobin variants ^48^. Alternatively, ADP-ribosylation could negatively affect the drug binding capabilities of plasma proteins, which would ultimately result in increased renal excretion of drugs, similar to what has been described for glycosylation of various drug binding plasma proteins ^50^. The data presented here suggest additional molecular insights into the function of plasma protein ADP-ribosylation. AHSG, which was modified in all 5 control pigs, but only 3 LPS pigs, is a secreted acute phase plasma protein whose concentration is routinely measured in clinical settings to roughly quantify ongoing inflammatory reactions^33, 51^. Examining the modification state of AHSG at different timepoints during infection, in addition to protein abundance, could potentially help further stratify disease bacterial infection duration and severity as well as potentially help predict clinical deterioration early-on.

Even though significant differences were not detected between the control and LPS groups, validating the modification of proteins uniquely modified in healthy or LPS-treated pigs could serve as potential diagnostic markers in a clinical setting. To this end, the detection of these low abundant proteins could be increased by depleting high abundant blood proteins^52^. Furthermore, the ADP-ribosylated proteins detected in plasma samples from LPS-treated but not healthy pigs could subsequently be validated as early biomarkers for SIRS. Many of the biomarkers currently validated for sepsis are not able to distinguish between infected and noninfected SIRS patients, and as a result, these diagnostic tests are not able to predict clinical deterioration^53^. Further development of the novel method described here could lead to the establishment of targeted-proteomic based diagnostics where plasma ADP-ribosylation patterns can be used to predict and prevent impending sepsis.

In conclusion, this study contributes to the rapidly developing field of ADP-ribosylation. To the best of our knowledge, this is the first study to define the *in vivo* whole blood or plasma ADP-ribosylomes. Importantly, the novel ADP-ribosylome workflow optimizations developed here can be used to define that WBL and plasma ADP-ribosylome of blood sample from any species, which could potentially advance our understanding of how this PTM contributed to disease development and manifestation within clinical settings. Indeed, further validation of the WBL and plasma ADP-ribosylome findings presented here could give rise to a vast array of new research possibilities and diagnostic tools. Finally, understanding the functions of extracellular protein ADP-ribosylation under healthy and disease conditions will help us unravel the underlying regulatory mechanisms of important cellular processes, like inflammatory responses, in the blood and further expand the functional processes associated with ADP-ribosylation.

## Acknowledgements

Dr. Peter Gehrig, Dr. Jonas Grossmann, and Laura Kunz (all Functional Genomics Center Zurich, University of Zurich) are kindly acknowledged for their support during MS analyses and Martina Lüthi (Institute of Plant Sciences, University of Bern) for her input on statistical analyses and proof-reading the manuscript. The animal experiment was funded by the Innosuisse - Swiss Innovation Agency as part of the CTI (2591.2 PFLS-LS). ADP-ribosylation research in the laboratory of MOH is funded by the Canton of Zurich and the Swiss National Science Foundation (grant 310030_176177).

## Author contributions

**Project conceptualization and administration:** M.O.H., D.M.L.P., S.C.L. (lead), R.G., X.S. (supporting, LPS trial)

**Investigation:** S.C.L., D.M.L.P. (lead); A.H., K.N. (supporting).

**Methodology:** S.C.L., D.M.L.P. (lead mass spectrometry and WB analyses) and A.H., K.N. (supporting); S.C.L., R.G. (lead animal handling and blood sampling), G.B., C.N. (production of LPS injections).

**Data curation and formal analysis:** S.C.L., D.M.L.P.

**Visualization and validation**: S.C.L., D.M.L.P.

**Writing, review & editing of MS**: S.C.L., D.M.L.P., M.O.H. (lead); K.N., R.G., X.S. (editing)

**Competing interests:** The authors declare no financial interest.

**Data and materials availability:** All data is available in the main text, the supplementary materials or in the ProteomeXchange Consortium database (PXDXXX).

**List of Supplementary Materials:**

- Suppl. Figure 1
- Suppl. Figure 2
- Suppl. Figure 3
- Suppl. Figure 4
- Suppl. Figure 5
- Suppl. Table 1
- Suppl. Table 2
- Suppl. Table 3
- Suppl. Table 4
- Suppl. Table 5
- Suppl. Table 6

