## Supplementary Information for "Development of a mass-spectrometry based method for the identification of the *in vivo* whole blood and plasma ADP-ribosylomes"

**Supplementary Table 1: Experimental groups, gender, weight and treatments of the pigs in the LPS trial.**

| Group | Gender | Weight (kg) | Treatment |
|---|---|---|---|
| control | F | 9.52 | NaCl |
| | M | 10.62 | NaCl |
| | M | 10.82 | NaCl |
| | F | 11.14 | NaCl |
| | M | 12.06 | NaCl |
| LPS | M | 10.48 | LPS |
| | F | 10.8 | LPS |
| | F | 10.86 | LPS |
| | M | 10.9 | LPS |
| | M | 11.2 | LPS |

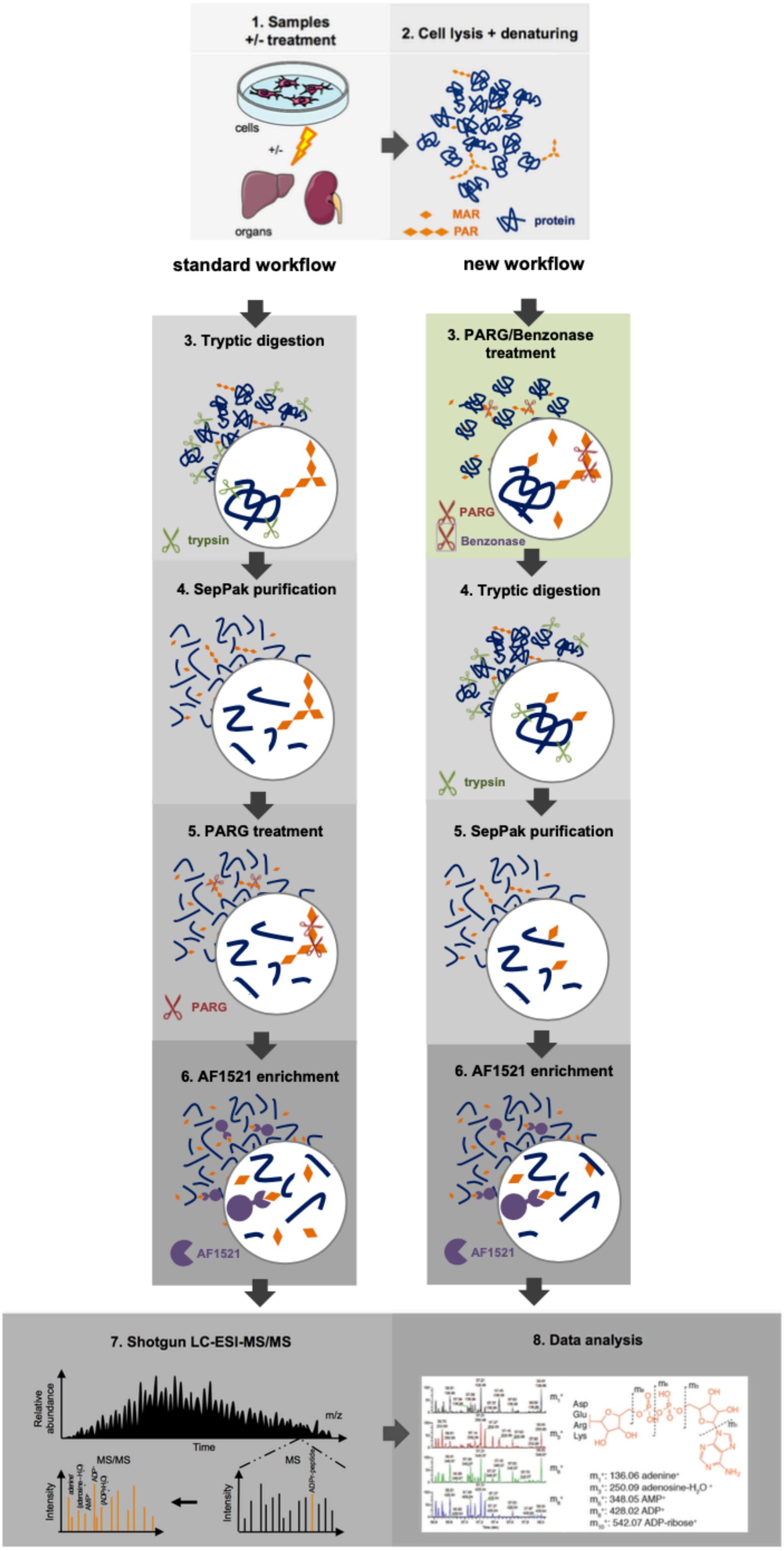


**Supplementary Figure 1: Overview of the standard workflow and new workflow for MS sample preparation.**

The standard workflow includes tryptic digestion of proteins, C18-SepPak peptide purification/clean-up, Af1521-based ADPr-modified peptide enrichment and a final pre-MS C18 stage-tip clean-up step (left column). For the new workflow, we moved the PARG/Benzonase treatment forward to the first step of the sample preparation protocol. This was done to optimize tryptic digestions and eliminate free ADP-ribose from the samples prior to Af1521-based ADPr-modified peptide enrichment (right column).

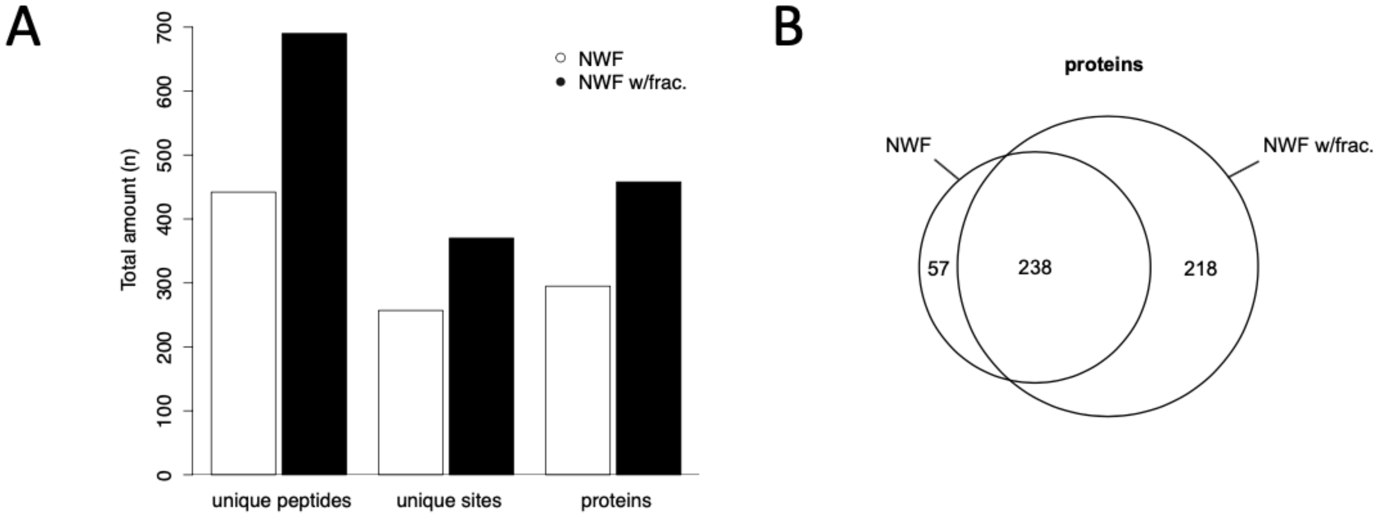


**Supplementary Figure 2: Comparison of the new workflow with and without LpH/HpH fractionation using HeLa $H_2O_2$-treated cell lysates.**

A: Comparison of the number of unique peptides, unique modification sites (>95% localization confidence) and proteins identified in HeLa $H_2O_2$-treated cell lysates when preparing the samples for LC-MS/MS using the new workflow with and without fractionation. Combined HCD and EthCD data is shown. B: Venn diagram showing the overlap of ADP-ribosylated proteins that were identified with the new workflow with and without fractionation.

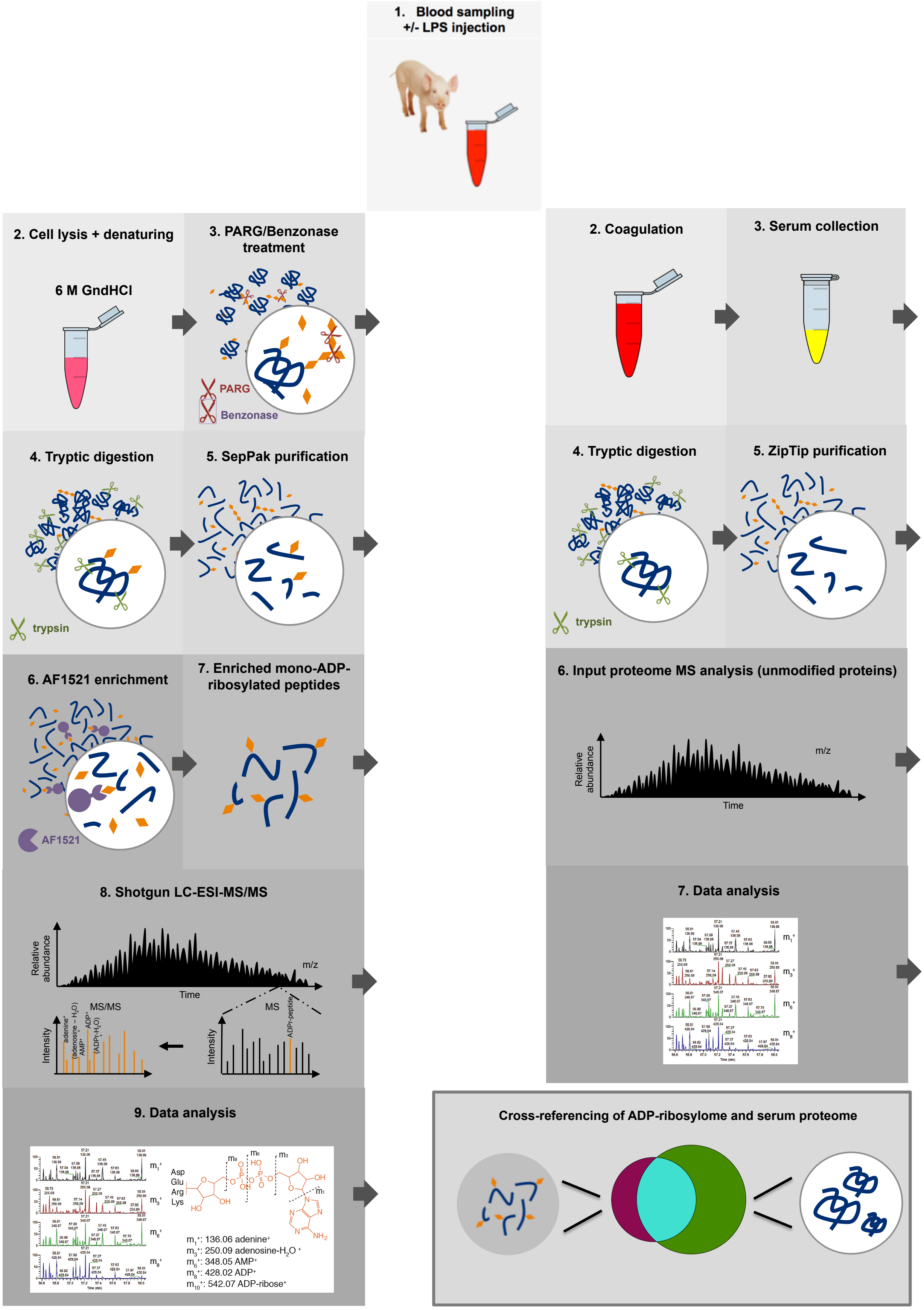


**Supplementary Figure 3: Overview of new ADP-ribosylome MS workflow used to identify the blood and plasma ADP-ribosylomes.**

ADP-ribosylated whole blood and plasma proteins were identified by applying the new workflow (left column) to WBL samples and comparing the modified proteins with the input plasma proteome (right panel).

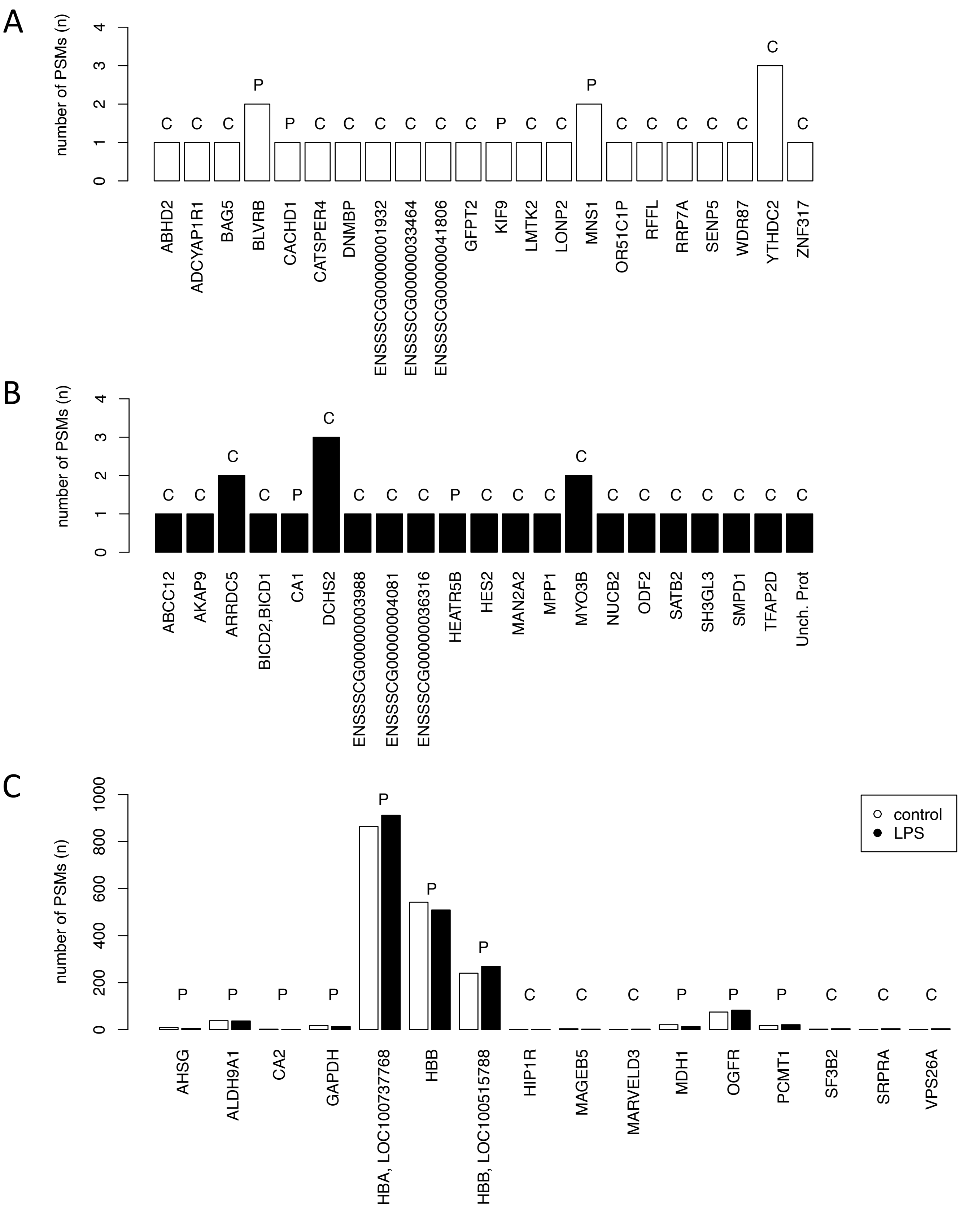


**Supplementary Figure 4: Unique, *in vivo* ADP-ribosylated proteins identified in healthy and LPS-treated pigs.**

A: Overview of the unique ADP-ribosylated proteins found in control pigs only. B: Overview of the unique ADP-ribosylated proteins found in LPS-treated pigs only C: Overview of the unique ADP-ribosylated proteins found in both control and LPS-treated pigs. “C” denotes proteins belonging to the cellular fraction of the blood and “P” those belonging to the plasma.
